# Functional effects of haemoglobin can be rescued by haptoglobin in an *in vitro* model of subarachnoid haemorrhage

**DOI:** 10.1101/2023.01.25.525148

**Authors:** Hannah Warming, Katrin Deinhardt, Patrick Garland, John More, Diederik Bulters, Ian Galea, Mariana Vargas-Caballero

## Abstract

During subarachnoid haemorrhage, a blood clot forms in the subarachnoid space releasing extracellular haemoglobin (Hb), which causes oxidative damage and cell death in surrounding tissues. High rates of disability and cognitive decline in SAH survivors is attributed to loss of neurons and functional connections during secondary brain injury. Haptoglobin sequesters Hb for clearance, but this scavenging system is overwhelmed after a haemorrhage. Whilst exogenous haptoglobin application can attenuate cytotoxicity of Hb and *in vivo*, and *in vivo* the functional effects of sub-lethal Hb concentrations on surviving neurons and whether cellular function can be protected with haptoglobin treatment remain unclear. Here we use cultured neurons to investigate neuronal health and function across a range of Hb concentrations to establish the thresholds for cellular damage and investigate synaptic function. Hb impairs ATP concentrations and cytoskeletal structure. At clinically relevant but sublethal Hb concentrations, synaptic AMPAR-driven currents are reduced, accompanied by a reduction in GluA1 subunit expression. Haptoglobin co-application can prevent these deficits by scavenging free Hb to reduce it to sub-threshold concentrations and does not need to be present at stoichiometric amounts to achieve efficacy. Haptoglobin itself does not impair measures of neuronal health and function at any concentration tested. Our data highlight a role for Hb in modifying synaptic function after SAH, which may link to impaired cognition or plasticity, and support the development of haptoglobin as a therapy for subarachnoid haemorrhage.

## Introduction

Subarachnoid haemorrhage (SAH) causes irreversible damage to brain tissues both during the acute phase and in secondary brain injury (SBI), leading to a high fatality rate of 30-40% [1,2] and significant disability in many survivors [3,4]. Raised intracranial pressure, oedema, inflammation and vasospasm lead to acute and delayed ischaemic damage, and are targeted by limited treatment options [4–7]. At the site of the haematoma, red blood cells (RBCs) begin to lyse within a few days, releasing their contents and leading to the accumulation of cell-free haemoglobin (Hb) in the cerebrospinal fluid (CSF). The Hb tetramer degrades into Hb dimers, hemichromes and eventually haem and free iron. Iron-containing products permeate into tissues, exposing neurons and other cells to oxidative stress [8]. Hb breakdown products such as met-Hb and haem can also directly activate inflammatory pathways, exacerbating damage and further contributing to SBI [9,10]. Hb and iron toxicity significantly contribute towards development of vasospasm and SBI [11–13], but are not currently clinically targeted.

It is well known that cell-free Hb leads to neuronal cell death *in vitro* [14–16] and *in vivo* [17,18] with key mediators being oxidative stress and ferroptosis caused by redox-active haem and iron, released from Hb [19,20]. More recently, research has suggested functional changes to neurons after exposure to Hb, such as reductions in synaptic anchoring proteins, neuroligins and neurexins [21] leading to reduced formation of excitatory synapses. The pre-synaptic marker synaptophysin and post-synaptic protein PSD-95 were downregulated in a mouse model of prolonged Hb exposure in ventricular and subarachnoid spaces [22]. In two rat blood injection models of SAH these biochemical changes were accompanied by cognitive deficits, indicating neurotransmission may be altered [23]. Further evidence shows that long-term potentiation (LTP), the synapse-strengthening process thought to underlie learning and memory [24], was impaired in a rat pre-chiasmatic injection model of SAH [25]. Hb-mediated impairments in synaptic plasticity as suggested by rodent studies may help explain poor rates of recovery seen in SAH, in addition to cognitive changes that affect quality of life in survivors [26,27]. Despite strong evidence linking Hb accumulation and iron deposition in the brain to neurodegeneration, both in SAH and a number of other neurodegenerative diseases [28–31], functional data on neuronal activity in the presence of Hb is limited.

Free Hb measured in the CSF after SAH is highly variable and peaks on average around 10 μM [13,22]. This peak typically occurs between day 10-12 after the onset thus presenting a wide therapeutic window for clinical intervention targeting Hb neurotoxicity. Haptoglobin (Hp) functions as an effective Hb scavenger by binding irreversibly to Hp and preventing haem release. Hp is typically present in blood plasma within the range of 0.3-3mg/ml [32– 34]. However, the large multimeric Hp proteins do not easily cross the blood-brain barrier and hence the concentration of Hp is much lower in the central nervous system compared to the systemic circulation [34], and Hp is rapidly depleted in the cerebrospinal fluid (CSF) after SAH [13]. Enhancing the CSF concentration of Hp *in vivo* can protect against cell loss, synaptic marker alteration, associated behavioural deficits [22] and vasospasm [12,22,35]. Furthermore, Hp administration has been used in clinical trials to prevent Hb-mediated kidney damage in sickle-cell anaemia and blood transfusion without significant adverse effects [36,37], indicating good peripheral tolerance.

Here we build on this knowledge by investigating the functional effects of persistent Hb exposure in neurons, measuring neuronal health and synaptic activity to understand the effects of clinically relevant Hb concentrations in the brain and how this may impact secondary brain injury after SAH. CSF analyses show that neurons experience extended exposure to 10 μM or more free Hb after SAH [13,22] and the composition of Hb species changes within the first days and weeks after SAH due to the conversion of Hb to met-Hb, further degradation, and dissociation into haem and free iron. This can be measured *in vivo* based on the paramagnetic properties of oxy-Hb, deoxy-Hb and met-Hb using magnetic resonance imaging (MRI), such that the composition of the blood clot appears distinctly different at SAH onset and in the days and weeks that follow [38,39]. Research has shown a delay in the onset of Hb-mediated cell death in neuronal culture, with cell loss starting 8 hours after application [14] suggesting that breakdown of Hb is likely implicated in its pathogenic effects. Whilst the exact timeline of Hb catabolism is uncertain in the brain environment, we chose to measure function of cells *vitro* after 48 hours and one week of Hb application, based on key changes to Hb composition assessed by neuroimaging.

We also studied the potential of Hp to prevent Hb-induced deficits, while considering the practical difficulties of the therapeutic agent reaching the location of haematoma and interstitial free Hb in a clinical setting. For example, is not likely possible to fully perfuse all CSF spaces with Hp after SAH, due to density of the haematoma in some places interfering with CSF flow [40]. High variability of free Hb concentration seen in patients after SAH will also be affected by the bleed volume, location and other factors [13], so accurately predicting Hb content and scavenging all free Hb is difficult. Additionally, not all free Hb is able to be bound by Hp *in vivo*: whilst the majority of Hb can be scavenged by Hp, there is a fraction of Hb that cannot be bound due to structural changes related to degradation or oxidative stress [22]. Hb permeates the outer cortex, as evidenced by an inward diminishing gradient of iron deposition after SAH [41], and it may be hard for a therapeutic agent delivered in the CSF space to reach the outer cortex in areas with closely apposed blood clot. Finally, the CSF is a purposely low protein fluid [42] and high CSF protein content is associated with poor outcome after SAH [43], which limits the amount of protein which can be administered intrathecally. For all these reasons, we investigated a partial, rather than stoichiometric, scavenging of free Hb by Hp, to understand the threshold of free Hb that is tolerated by neurons. Ultimately this will improve understanding of the effects of Hb on neuronal function and add to the body of knowledge needed to develop Hp as a clinical therapeutic in SAH.

## Materials and Methods

### Haemolysate preparation

Human blood was obtained from human volunteers after informed consent (National Research Ethics Service approval 11/SC/0204 and institutional research ethics approval ERGO 41084.A1). Whole blood was collected in heparin tubes (BD, UK) and transferred to a centrifuge tube onto a layer of histopaque-1077 (Sigma-Aldrich, roughly 30% of final volume). The tube was centrifuged at 900G for 15 minutes and top layers removed, leaving the lower fraction containing red blood cells (RBCs). RBCs were washed with sterile Dulbecco’s phosphate buffered saline to remove other cells and plasma until supernatant was clear and colourless, before lysing with sterile distilled water. Following centrifugation at 13,000rpm for 30 minutes to remove ghost membranes, supernatant was removed, passed through a 15 μm filter and analysed with a bicinchoninic acid assay (Pierce™□) to determine protein concentration before storage at -80°C. Spectrophotometry was carried out on each batch before use to determine the haemoglobin species composition, comprising on average 79.8 ± 1.4% oxyhaemoglobin, 13.2 ± 1.3% deoxyhaemoglobin and 7.0 ± 2.0% methaemoglobin [44]. Haemolysate (HL) concentration is expressed throughout as a concentration of dimers, hence 10 μM HL is equivalent to 20 μM of iron-containing haem.

### Source of haptoglobin and scavenging analyses

Hp was prepared by Bio Products Laboratory Ltd from pooled human blood plasma, enriched for Hp1 dimer. Hp was dialysed with a molecular cut off of 14 kDa and endotoxin content was measured at <0.02 E.U./ml. Hp molar concentration refers to a weighted average molecular weight of monomers, in the purified mixture of Hp1 and Hp2.

To determine rate of free Hb scavenging, a fixed amount of HL was incubated with increasing amounts of Hp for 16 hours at room temperature, and 2 μg of Hb from each mix separated on a non-denaturing 8% polyacrylamide gel. Hb band density was analysed against a HL-only control lane within each gel to quantify the percentage of unbound Hb for each incubation ratio. A binding curve from three gel repeats was used to estimate the concentration of free Hb in treatment groups with co-application of Hp throughout experiments.

### Primary neuron cultures

C57/BL6 mice of either sex (Charles River, bred in-house under a 12/12 hr light/dark cycle at 21°C) were culled on the day of birth (P0) and brains dissected in Dulbecco’s PBS without calcium or magnesium (Gibco). Hippocampi were isolated and dissociated with papain before seeding onto poly-D-lysine coated glass coverslips at 1,000 cells/mm. Cells were incubated with Neurobasal Medium (Gibco) supplemented with 1% Glutamax-I and 2% B27, incubated at 37°C with 5% CO_2_. A full media change was carried out at DIV7 and HL and/or Hp applied after randomised allocation to culture wells alongside a media volume top-up at DIV14. Treatment groups were blinded until raw data analysis was complete for each experiment.

## ATP Assay

Primary cultures were washed once with supplemented Neurobasal Medium and then fresh medium applied, with an equal volume of reaction mixture from the CellTitre Glo assay kit (Promega) at room temperature to measure adenosine triphosphate (ATP) levels. The kit was used as per manufacturer instructions: plates were mixed using an orbital shaker for 2 minutes, incubated for a further 8 minutes before 200 μl of mixture was pipetted in triplicates into 96-well plates for measurement of luminescence, using a Promega Glo-Max®-Multi Microplate reader. Background luminescence was measured and subtracted from readings using a control of culture medium without cells, and results were unblinded and normalised to vehicle-treated cells within cultures.

### Immunofluorescent staining and microscopy

Neuron cultures were fixed at DIV21 with 4% paraformaldehyde, permeabilised with 0.1% Triton-X100 and washed in tris-buffered saline. Cells were blocked using goat serum and stained with anti-tubulin 1:400 (Cell Signalling 2128) followed by Alexa Fluor secondary antibodies at 1:1000 in 2% goat serum. Coverslips were mounted onto microscope slides using mounting medium containing 4⍰,6-diamidino-2-phenylindole (DAPI) (Fluoroshield ab104139, Abcam) and imaged using an Olympus IX83 inverted microscope with a 40X air objective, Intensilight CHGFI metal halide light source (Nikon) and Optimos sCMOS camera (photometrics, USA). Images were acquired using Cellsens software and processed by background subtraction using a radius of 50 pixels and overlaid using FIJI software [45]. Treatment groups were blinded from the point of treatment application until image processing was completed, and fields of view were chosen using DAPI channel.

### Patch clamp electrophysiology

Cells were used for patch clamp electrophysiology at DIV21 ± 1. Recordings were low-pass filtered at 5 kHz using an Axopatch 200B amplifier and acquired at 20 kHz using a National Instruments board analog to digital converter. Matlab software (Mathworks, Natick, U.S.A.), custom software (Ginj2.0, Hugh P.C. Robinson) and WinEDR software (Strathclyde) were used for data acquisition. Borosilicate glass micropipettes were pulled to a resistance of 5-7 MΩ and filled with intracellular solution containing, in mM, 125 potassium gluconate, 10 KCl, 10 HEPES, 10 Phosphocreatine, 0.4 GTP, 4 Mg-ATP and pH balanced to 7.3 using KOH. Recordings were made in artificial CSF containing, in mM, 126 sodium chloride, 2 calcium dichloride, 10 glucose, 2 MgSO4, 3 potassium chloride and 26.4 sodium carbonate. The liquid junction potential of -12.5 mV was not corrected for. Artificial CSF was bubbled with 95% oxygen, 5% CO2 throughout and maintained at 25 ± 1°C.

Passive membrane properties were measured using voltage-clamp, and cells excluded if the input resistance exceeded 1 GΩ indicating non-pyramidal cells [46–48], or if series resistance exceeded 30 MΩ. The first 3 recorded cells to meet these criteria in each repeat were included for further analysis. Active membrane properties were measured using a current-step injection stimulus in bridge-compensated current clamp (see Figure 3), and current injection to maintain cells at -70 mV. Evoked EPSPs were recorded from connected pairs of cells. A positive current injection of 500-1000 pA with duration 6 msec was applied to induce a single action potential and measure a postsynaptic response repeated at 0.14 Hz. Evoked excitatory postsynaptic potentials (EPSCs) were recorded in the same manner, with the postsynaptic cell in bridge mode. Paired-pulse ratio was measured from EPSCs with a 50 msec interval between presynaptic stimulations. Miniature EPSCs (mEPSCs) were recorded at -70 mV using a Cs-gluconate based intracellular solution containing, in mM, gluconic acid 70, caesium chloride 10, sodium chloride 5, BAPTA 10, HEPES 10, QX-314 10, GTP-NaCl 0.3, ATP-Mg 4 and adjusted to pH 7.3 using 1M CsOH. Artificial CSF contained 500 nM tetrodotoxin and 1 μM SR-95531. The liquid junction potential for Cs-gluconate solution of -12 mV was not corrected for.

### Polyacrylamide gel electrophoresis

Protein lysates were collected at DIV21 for Western blotting and separated in 7% sodium dodecyl sulfate polyacrylamide gels using electrophoresis. Proteins were transferred to nitrocellulose membranes and stained using anti-GluA1 antibody (13185, Cell Signalling) secondary antibody & imaged using an infrared scanner (Odyssey®, LI-COR Biosciences). Blot quantification was carried out using ImageStudio software (LI-COR). For Hb-Hp binding experiments, a fixed amount of Hb was incubated with varying Hp at room temperature overnight and the resulting mixtures separated using non-denaturing 8% polyacrylamide gel electrophoresis before staining with Coomassie, loading 2 μg of Hb per lane. Gels were de-stained in 10% methanol with 7% glacial acetic acid overnight, imaged and quantified against a control lane containing 2 μg of unbound Hb within each gel.

### Data analysis

mEPSC recordings were analysed with Eventer software (A. Penn, University of Sussex [49]) with a low-pass filter of 1000 Hz. A machine-learning model was trained using approximately 2500 mEPSC events across all treatment conditions, detected using the Pearson method with a threshold of 4 standard deviations of the noise. Training took place by manually accepting or rejecting events based on appropriate rise and decay kinetics. The model was then employed to detect events automatically in each mEPSC recording and report the inter-event interval and peak amplitude of the first 500 events per cell. Intrinsic membrane properties and evoked EPSP and EPSC amplitude were analysed from raw data using custom-written Matlab code. Evoked currents and potentials were averaged from 50-100 repeats per cell, 3 cells per independent neuron culture with automatic event onset detection. Paired-pulse ratio was measured as the ratio between median average values for each EPSC from 50-100 repeats per cell. All statistical analyses were performed in GraphPad Prism V9 (GraphPad Software, CA, USA) using a two-way repeated measures ANOVA with matching within cultures and Dunnett’s post-hoc corrections. We did not assume data sphericity in experiments where data was normalised within each culture, and epsilon (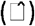) is reported when the Geisser-Greenhouse correction has been applied in these analyses. Data are represented as mean ± SEM.

## Results

### Haemolysate disrupts ATP levels and neurite structure *in vitro*

To firstly characterise the threshold for neuronal damage by Hb in our neuronal culture system we applied increasing concentrations of HL prepared from human RBCs to hippocampal neuronal cultures at DIV14 and we measured ATP concentration in the lysed cell population using the luciferase-based CellTitre Glo assay. HL application started at 10 μM of Hb dimer as measured in human CSF after SAH [22], and was applied up to 200 μM. We observed a dose-dependent reduction in ATP starting at 50 μM HL at 48 hours (F(2.4,23.6)=520.8, 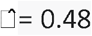, p<0.0001, Figure 1A), and starting at 20 μM after one week of HL application (F(2.5,29.9)=713.6, 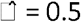, p<0.0001, Figure 1B). To assess neurite structure, we imaged neurons using immunofluorescent staining of β-tubulin and DAPI. At all concentrations of HL used, neurite beading indicative of microtubule structure disruption was observed after a one week incubation, which worsened with increasing concentration of HL (Figure 1C).

**Figure 1.**
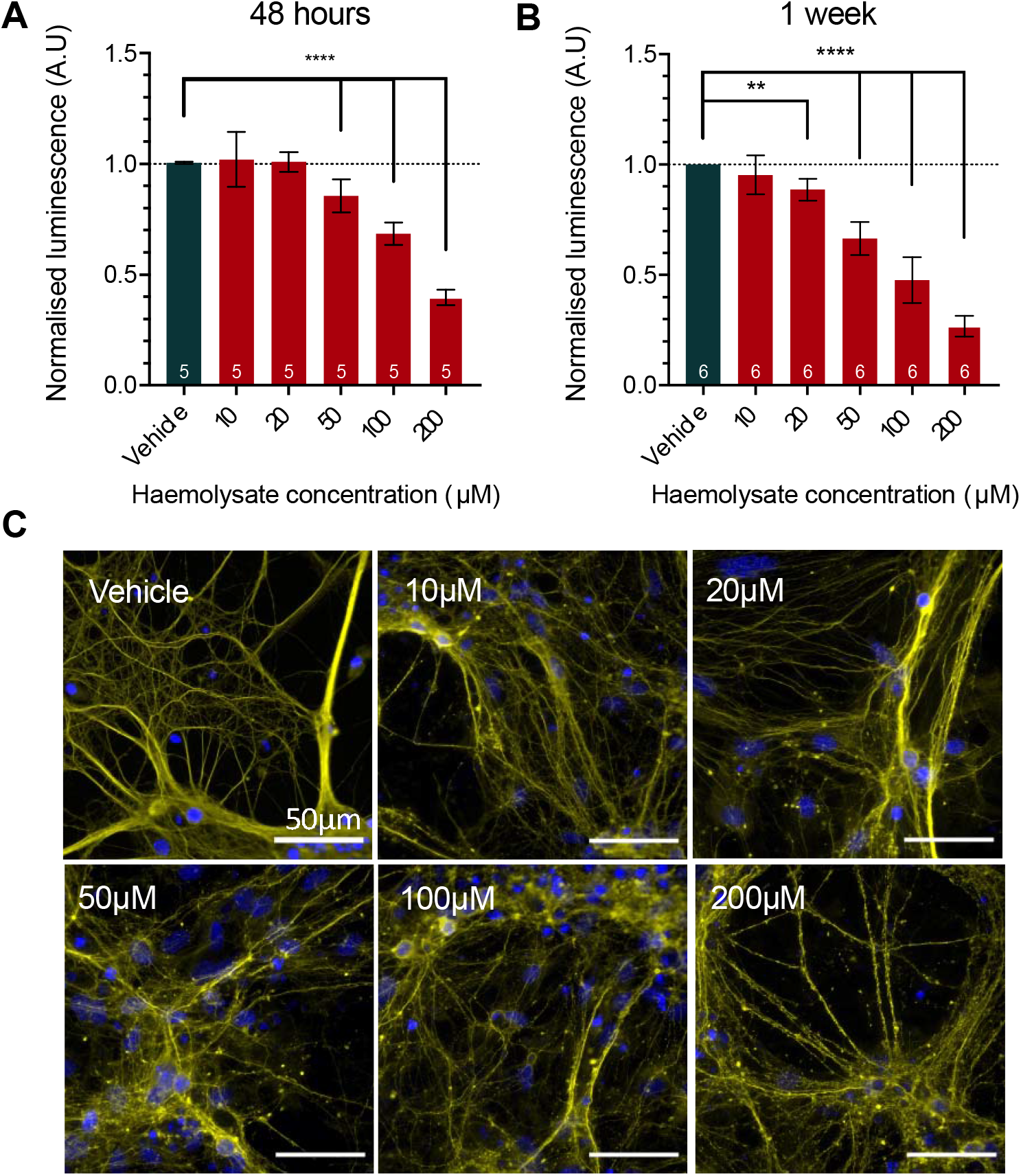
Haemolysate impairs ATP levels and neurites in cultured neurons. Primary hippocampal neurons were incubated with haemolysate from DIV14. A) The CellTitre Glo assay found a reduction in ATP concentration after 48 hours and B) one week of exposure to haemolysate. C) Immunofluorescent staining of β tubulin shows disruption of cytoskeletal microtubules as neurite beading in the presence of haemolysate after one week. Significance levels: ** = P<0.01, **** P<0.0001.

### Haemolysate neurotoxicity can be prevented with haptoglobin

To determine if Hp can prevent HL-induced toxicity, we co-applied Hp with HL to cells for one week and repeated ATP measurements. Hp alone at 30 μM appeared to show an increase in ATP (F(2.58,25.8)=40.4, 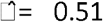, p<0.0001, Figure 2A) but all other concentrations up to 120 μM were not different compared to vehicle conditions indicating that Hp does not reduce ATP levels in neuronal cell cultures even at very high concentrations.

**Figure 2.**
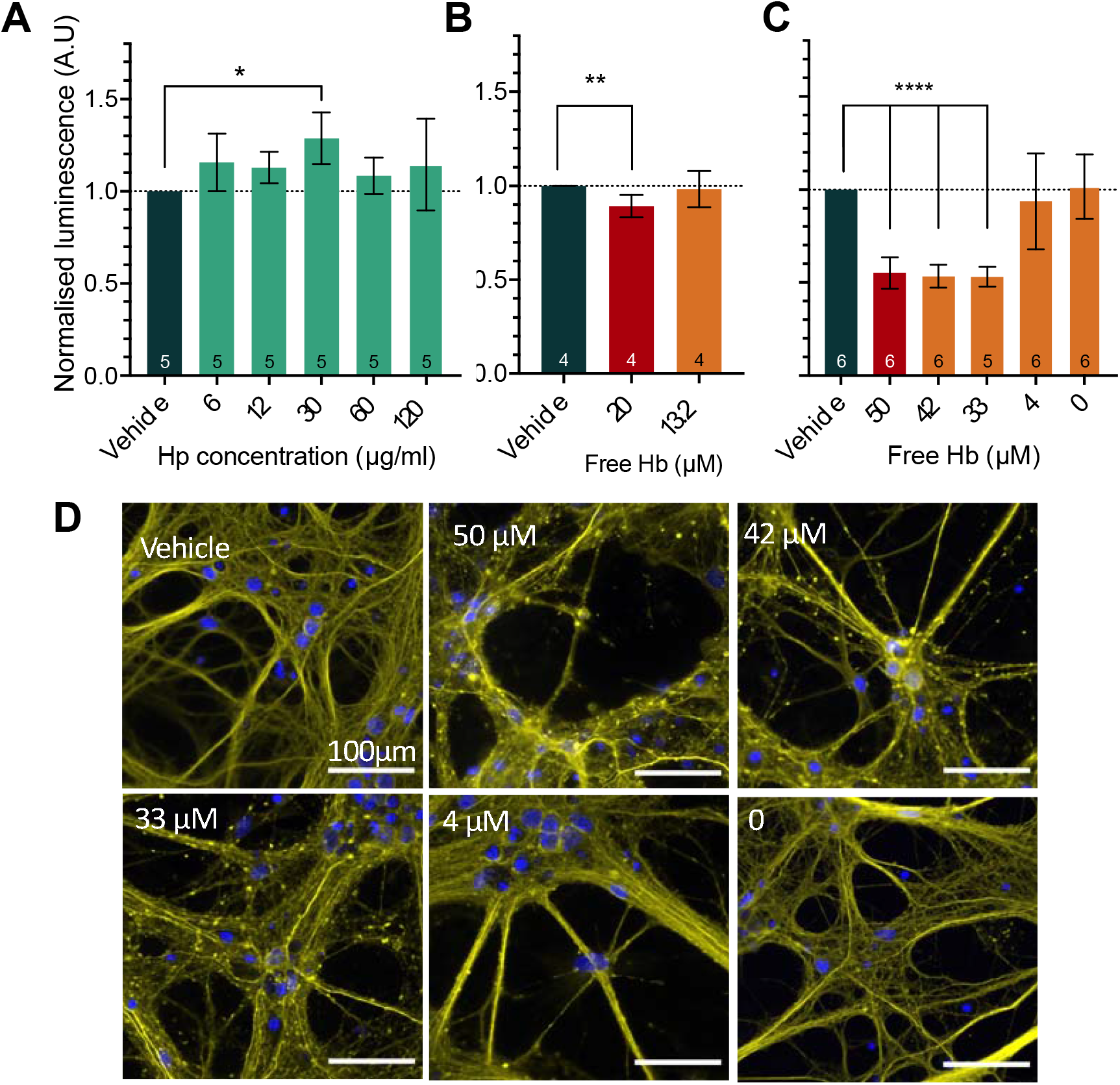
Haptoglobin can prevent deficits in ATP and neurite structure caused by haemolysate. A) ATP concentration in cultured neurons after one week of exposure to haptoglobin up to 120 μM. B) ATP levels after incubation with 20 μM of HL (red bars) and co-application of Hp to scavenge one-third of free Hb (orange). C) 50 μM HL co-applied with increasing amounts of Hp to scavenge free Hb. D) β-tubulin staining shows disrupted microtubule morphology after incubation with 50 μM HL and co-application of Hp, expressed as concentration of free Hb remaining in media. Significance levels: * = p<0.05, ** = p<0.01, **** p<0.0001.

**Figure 3.**
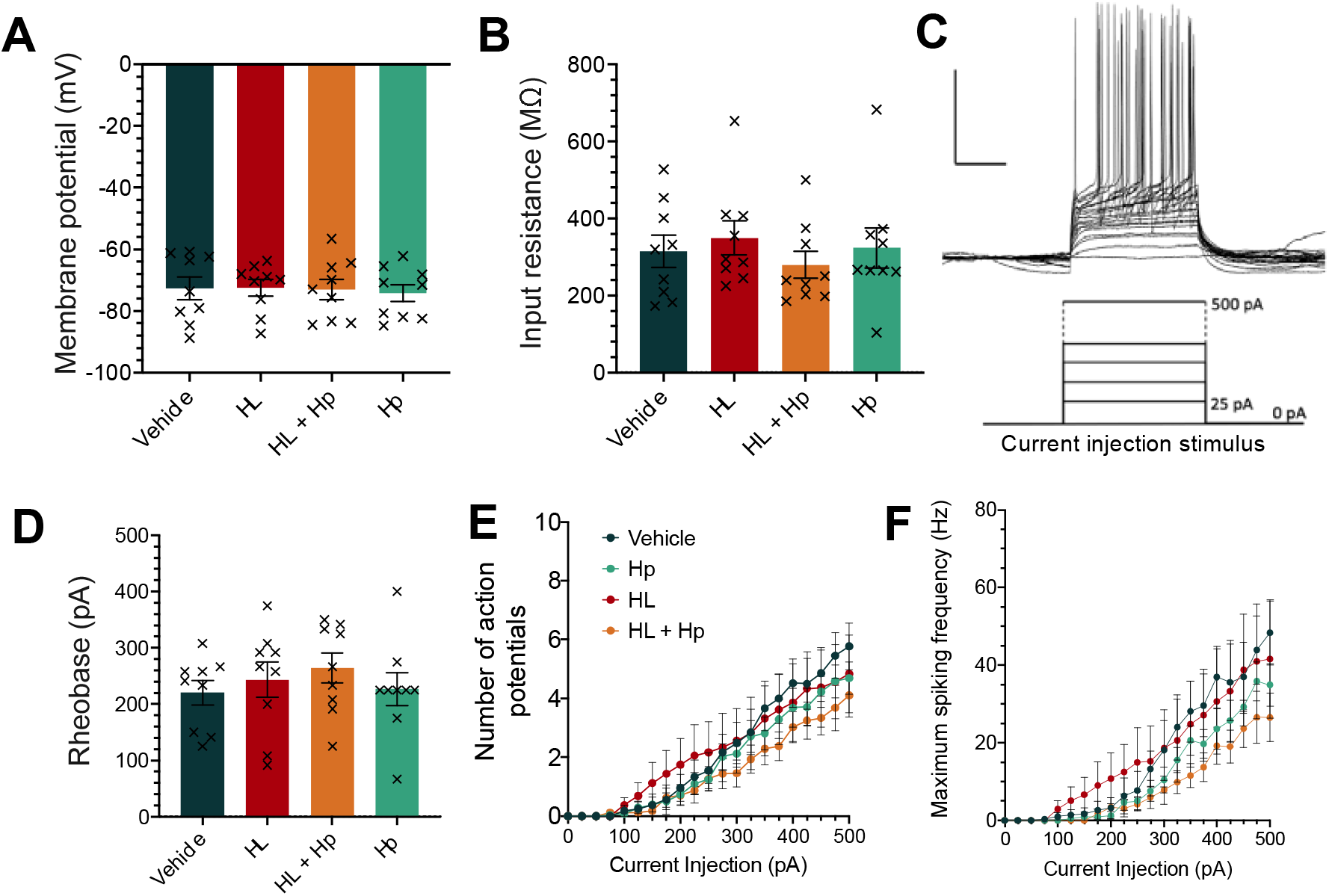
Intrinsic neuronal membrane properties are not altered by 10 μM Hb within one week. A) Membrane potential and B) input resistance are measured upon break-in to the cell in whole-cell patch clamp. C) A current injection stimulus in steps of 25 pA is applied to the cell in bridge compensated current clamp. D) Rheobase measured from a 250 msec current injection in C. E) maximum number of action potentials fired and F) the maximum spiking frequency observed at each current injection step. N = 3-5 cells per culture, 9 cultures.

Next, we investigated whether partial scavenging was sufficient to protect from neurotoxicity by binding one-third of free Hb with exogenously co-applied Hp. We measured ATP levels after co-applying Hp with HL for one week, and found that scavenging 34 ± 3.6 % of free Hb, reducing free Hb from 20 to 13.2 ± 0.7 μM, was sufficient to prevent an ATP deficit (F(1.2, 9.7)=12.4, 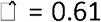, p=0.004, Figure 2B). We further investigated 50 μM HL with increasing Hp concentrations, and found that the ATP deficit was not prevented by scavenging one-third of free Hb, but could only be prevented when the majority of free Hb was bound by Hp (F(2.2,43.8)=124.9,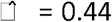, p<0.0001, Figure 2C). These data suggest that Hp can prevent ATP deficits in neuron cultures at high concentrations of Hb, by scavenging free Hb to sub-lethal levels. The threshold for a Hb-induced ATP deficit appeared to lie between 13.2-20 μM free Hb. Staining of β-tubulin in the same neuronal cultures after incubation with 50 μM HL and co-application of Hp showed neurite beading in all conditions with free Hb, and microtubules were restored to vehicle morphology under conditions of full Hb scavenging (Figure 2D).

### Sublethal haemolysate does not alter intrinsic membrane properties of neurons

Previous data had shown that free Hb perturbs the resting membrane potential of neurons [50] and these effects may be driven by changes in neuronal excitability or ongoing synaptic activity. To investigate the functional effects of HL on the excitability of neuronal cultures we measured intrinsic membrane properties of neurons at DIV21 ±1 in whole-cell patch clamp electrophysiology, after a one week exposure to HL or Hp in culture medium. 10 μM Hb, which has been measured as the average peak in human CSF after SAH [22] does not appear to cause an ATP deficit and was therefore used as a clinically relevant yet sublethal HL insult for electrophysiological studies. As such we could use visualised patch clamp to record from neurons randomly selected from a large population. Since scavenging one-third of free Hb restored ATP deficits using 20 μM HL, we employed the same scavenging ratio when co-applying Hp to 20 μM HL for electrophysiology, reducing the free Hb concentration in culture medium from 10 μM to approximately 6.4 ± 0.4 μM.

We found no effect of treatment on the resting membrane potential of neurons (F(3, 54)=0.2451, p=0.86) or input resistance (F(3, 54)=1.34, p=0.27 (Figure 3A-B). Next, we used current clamp to apply a current injection stimulus to the cells and analysed the rheobase (F(3, 54)=1.08, p=0.36, Figure 3C-D). Finally, we quantified the number of action potentials fired at each current injection step (Figure 3E), and there was no effect of treatment (action potential number: F(3, 15)=0.52, p=0.68, maximum frequency: F(3, 15)=0.85, p=0.49). The same membrane properties were also measured at 48 hours of exposure to HL or Hp, with no effect of treatment across all analyses (data not shown).

Our data indicate that cultured neurons can maintain their intrinsic membrane properties throughout a one-week exposure to 10 μM HL. This corroborates the finding that there is no disturbance to ATP levels in these cultures at this concentration.

### 10 μM HL reduces AMPA receptor-mediated synaptic currents

We investigated synaptic function in neurons exposed to HL by measuring mEPSCs. These currents represent the activation of α-amino-3-hydroxy-5-methyl-4-isoxazolepropionic acid (AMPA) receptors due to spontaneous fusion of presynaptic vesicles and glutamate release. There was no change in frequency of mEPSCs (treatment effect F(3,54)=1.98, p=0.13, Figure 4B), but we observed a reduction in amplitude in the presence of HL compared to vehicle (F(3,54)=4.99, p=0.004, HL vs vehicle p=0.001, Figure 4C) which was not observed with Hp co-application (p=0.18, Figure 4C).

**Figure 4.**
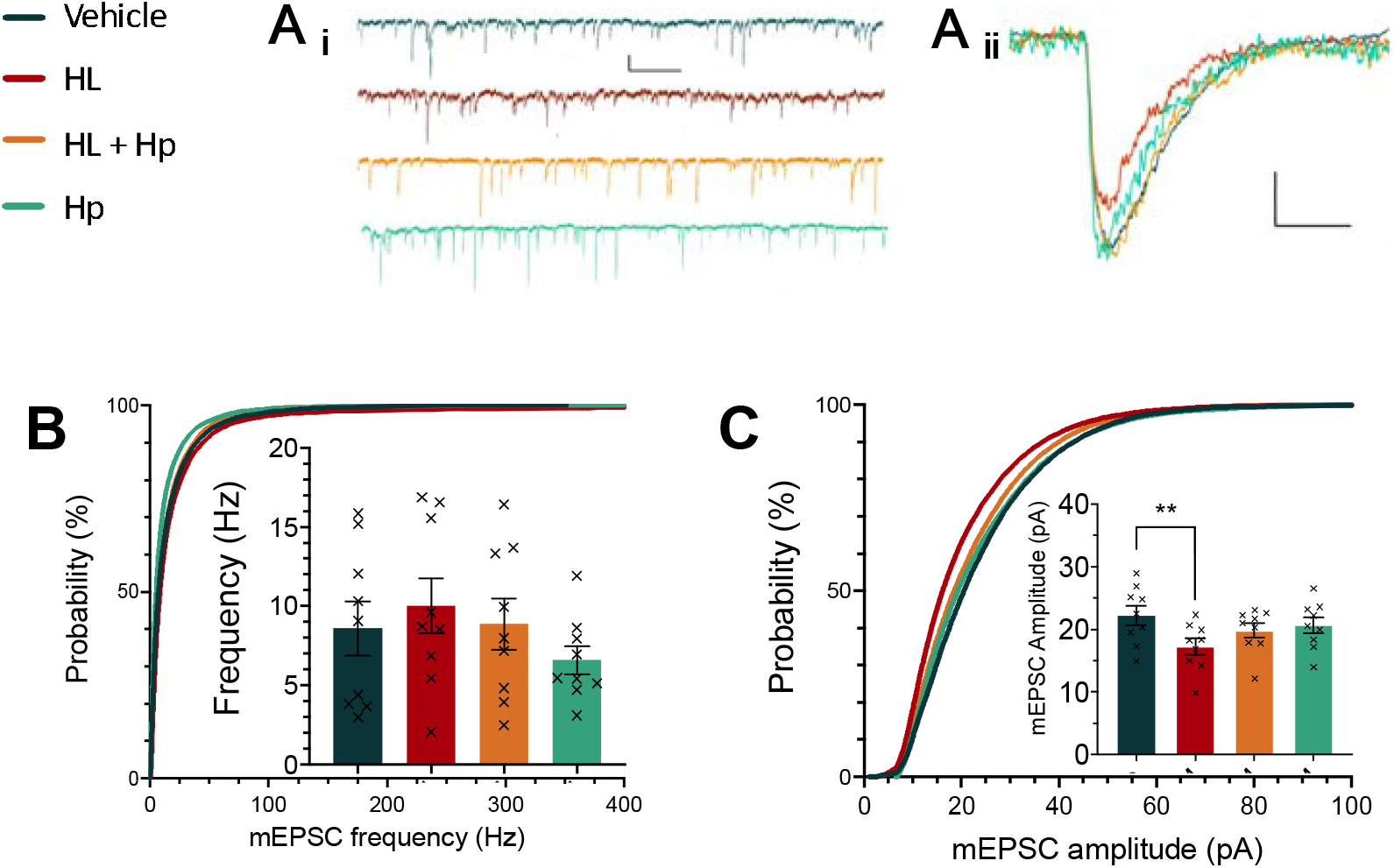
Miniature excitatory postsynaptic current (mEPSC) amplitude is reduced by a 1-week exposure to 10 μM HL. A) Sample traces showing i) mEPSCs (scale bar 1 sec/10 pA) and ii) overlaid individual events (scale bar 10 msec/ 10 pA). B) Quantification of mEPSC frequency and C) amplitude as cumulative frequency distribution and median value per culture. N = 3 cells per culture, 9 cultures. Significance levels: ** p<0.01

Next, we quantified unitary EPSPs in connected cell pairs. After a one week exposure to 10 μM HL, the EPSP amplitude was significantly decreased (F(3,24)=3.54, p=0.03, HL vs vehicle p=0.01, Figure 4D-E) and this effect was prevented when Hp was co-applied to reduce free Hb (vehicle vs HL + Hp: p=0.15). We also quantified the unitary EPSC, which showed a similar reduction in amplitude in the presence of HL (F(3,24)=3.41, p=0.034, HL vs vehicle p=0.015) but not when Hp was also present (p=0.16, Figure 4G). The paired-pulse ratio was not different across conditions (F(3,24)=0.48, p=0.70), nor was the failure rate (vehicle: 3.39 ± 2.2 %, HL 3.5 ± 2.7 %; treatment effect F(3,24)=1.57, p=0.22).

When HL and Hp were co-applied, AMPAR currents were not significantly different to HL alone, but were not different from vehicle conditions either. This indicates Hp was providing a partial rescue of synaptic AMPAR currents. To ensure Hp was not suppressing AMPAR-mediated currents, we measured evoked EPSP and EPSC amplitude at a higher concentration of 24 μM Hp and found no difference to vehicle (EPSP amplitude: vehicle 1.93 ± 0.34 mV, Hp 24 μM 2.47 ± 0.59 mV, p=0.48. EPSC current: vehicle 42.6 ± 11.7 pA, Hp 24 μM 41.3 ±15.2 pA, p=0.95. N = 3 pairs).

### 10μM haemolysate reduces GluA1 expression

To assess if the protein levels of one of the major subunit components of AMPA receptors, GluA1, is affected by HL exposure we prepared protein lysates from cell cultures at DIV21 after one week of incubation with 10 μM HL with or without Hp, and separated using SDS-PAGE and Western blotting. Membranes were probed for GluA1 and normalised to β-tubulin. We found a reduction in GluA1 expression in cells exposed to HL (F(3,12)=5.12, p< 0.05, HL Vs Vehicle p<0.05) but not when Hp was co-applied (p=0.20, Figure 5B) indicating downregulation of GluA1 protein in the presence of unbound Hb.

**Figure 5.**
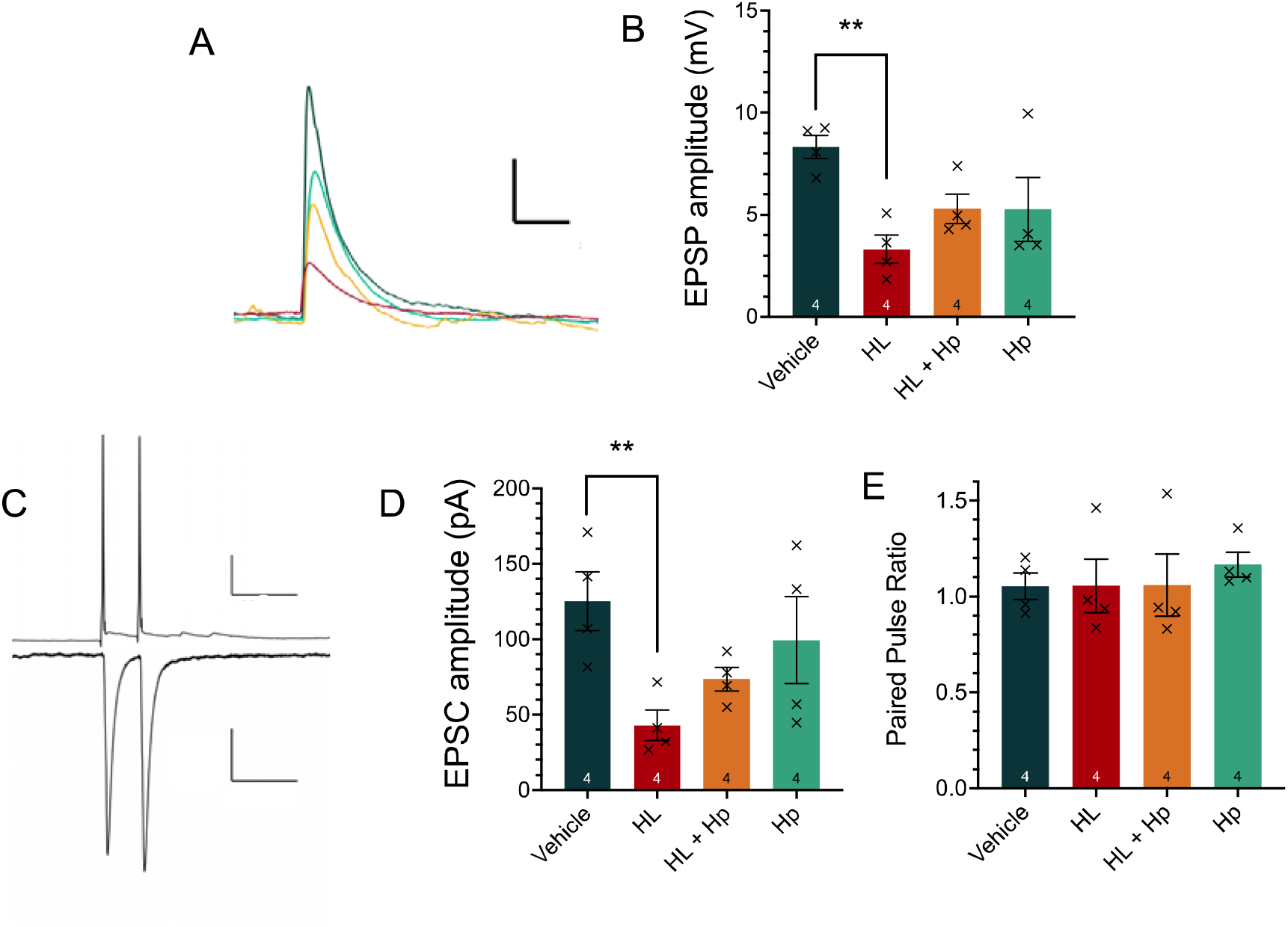
Amplitude of evoked excitatory post synaptic potentials (EPSPs) and excitatory postsynaptic currents (EPSCs) is reduced by 10 μM HL. A) Representative mean EPSPs (scale bar 50 sec/ 2 mV). B) Quantification of evoked EPSP amplitude. C) Paired-pulse sample trace with inter-pulse interval of 50 msec (scale bar top: 100 msec/ 25 mV, bottom: 100 msec/ 25 pA). D) Unitary EPSC amplitude and E) Paired pulse ratio. N = 3 pairs per culture, 4 cultures. Significance levels: ** p<0.01.

**Figure 6.**
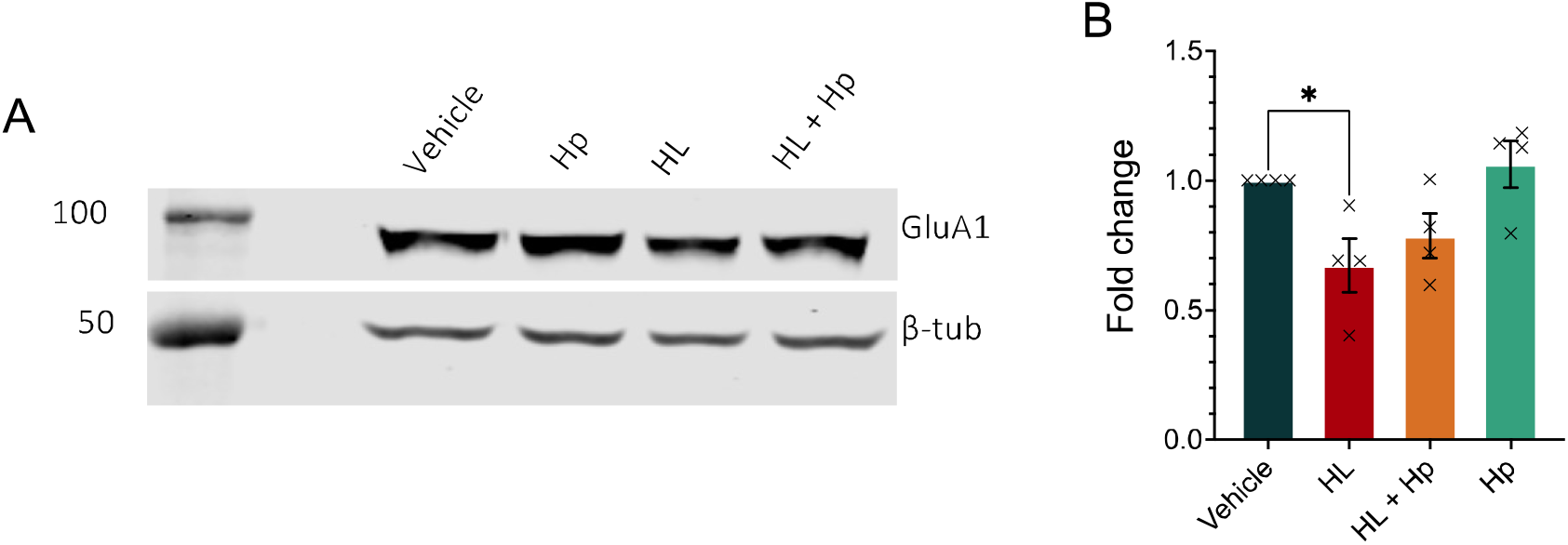
Total AMPA receptor GluR1 subunit levels are reduced in hippocampal cell cultures by a one week incubation with HL. A) Hippocampal cell culture lysates were prepared at DIV21 after a one week incubation with 10 μM HL or Hp and blots probed for GluR1 and β-tubulin. B) Quantification of GluA1 normalised to loading control. N = 3 cultures. Significance level: * p<0.05

## Discussion

We investigated the effects of Hb at concentrations relevant for SAH on neuronal health and synaptic activity and determined whether partially scavenging free Hb can prevent functional deficits in neurons. SAH features a complex variety of pathological mechanisms contributing to acute damage and secondary brain injury, which have been extensively reviewed [51–54]. Hb accumulation in the subarachnoid space through haemolysis exposes neurons to an average peak of 10 μM Hb in CSF, and possibly higher concentrations at the site of the haematoma. This Hb peak occurs at between 10-12 days after SAH onset, as seen in human CSF analyses [13,22]. CSF concentrations of Hb have been proposed as a diagnostic marker for SBI, with a proposed threshold of 7.1 μM of tetrameric Hb [13], equivalent to 14.2 μM of Hb dimer, hence similar to the present study. In another study, the average Hb peak concentration in CSF was higher in patients who developed delayed ischaemic neurological deficits (10μM) than those who did not (6 μM) [55]. This evidence suggests that the level of CSF Hb is directly linked to SBI and hence poor recovery after SAH. Scavenging Hb to reduce its concentration in the brain has potential as a therapeutic intervention, and there is generous timeframe to intervene.

To first characterise the threshold of free Hb that causes damage within our neuronal culture model, we measured ATP levels in the presence of Hb for up to one week. We found that the threshold for damage measured as an ATP deficit at 48 hours (50 μM) was higher than that at one week (20 μM), suggesting Hb neurotoxicity occurs progressively as previously suggested [16] and impairs ATP levels. This may occur through suppression of metabolic processes or loss of cells. Primary hippocampal cultures contain glial cells unless treated with cell cycle inhibitors [56,57], which provide trophic support and may contribute to ATP levels in these assays. However, evidence suggests that neurons are more vulnerable to Hb-mediated cell loss than glia, as in co-culture models glial cells and oligodendrocytes appear unaffected by Hb exposure at similar concentrations [14,15,58]. We cannot exclude effects of HL on glia, or of glial cell contribution to ATP levels and therefore to identify neuron-specific effects of Hb we immunostained cultured cells for β-tubulin to identify neurites. We observed neurite beading, indicative of cytoskeletal breakdown such as that seen in Wallerian degeneration [59] that worsened with increasing concentration of Hb. This occurred even at 10 μM free Hb, where an ATP deficit was not observed, suggesting that microtubule degradation may occur prior to loss of cells or metabolic deficit. Along the same lines, degeneration of axons was seen after SAH in humans, as reflected by elevations in CSF neurofilament-light level [22,60,61], long considered a key marker of neurodegeneration, and this correlated with preceding CSF Hb concentration [22,62].

We next investigated the potential of purified human Hp to protect from Hb-mediated damage, by co-incubating cell cultures with HL and Hp for one week. The differential efficacy of Hp genotypes in scavenging Hb from the brain and outcome after SAH has been reviewed previously [63–65] with a general theme that the presence of Hp1 is beneficial. Hp1-1 and Hp1-2 genotypes cluster closely as having better neuroprotective qualities than Hp2-2. In this study we have used Hp from pooled human blood plasma, containing both isoforms but enriched for Hp1 protein, containing approximately 60% Hp1-1 dimer.

We considered the logistics of Hp infusion into the subarachnoid space containing a peak CSF concentration of 10 μM Hb. Under these conditions, if one aims to bind all free Hb based on an average molecular weight for Hp monomers of 52.18 kDa within the Hp preparation used in this study, a CSF Hp concentration of 0.522g/L would need to be achieved. Protein levels in the CSF in healthy individuals vary in the range 0.3-0.4 g/L [66,67], increasing to 0.7-0.8 g/L after SAH [68]. There is an association between high CSF protein and poor outcome after SAH, linked to higher rates of SBI [43]. We therefore aimed to model a sub-stoichiometric, or partial scavenge, of free Hb *in vitro*, to recapitulate the likely situation in vivo more realistically. As discussed in the introduction, a partial scavenge is more likely to be achievable in a clinical setting due to variable factors in bleed volume and hence Hb concentration, density of the haematoma and modifications to Hb over time making it unable to be bound by Hp [13,22,40].

One previous report suggests that Hp (also from Bio Products Laboratory) can enhance Hb-mediated neuronal cell loss [69], whilst another study showed that the same Hp formulation can prevent Hb toxicity in cultured neurons [22]. To investigate the possibility of Hp being toxic in itself, we tested Hp alone up to high concentration of 120 μM and found no impairment of ATP levels, in fact a small increase in ATP was found when 30 μM Hp was applied for one week. After applying HL with or without Hp, we found that ATP levels were restored to vehicle once free Hb was reduced to 13.2 ± 0.7 μM or less, regardless of whether 20 or 50 μM HL had been applied. In microtubule microscopy, complete binding of all free Hb showed a full restoration of neurite integrity, so that even 50 μM of Hb once in complex with Hp was no longer damaging to the cells. In conclusion, free Hb must be reduced to sub-lethal levels of between 13.2-20 μM to prevent neuronal damage as indicated by ATP deficits within our culture system, and this can be achieved using Hp. Furthermore, Hp prevents Hb-induced ATP or microtubule impairment. Our results agree with the recent publication by Garland and colleagues demonstrating that Hp can prevent, rather than enhance, Hb-mediated toxicity [22].

Next, we measured the function of neurons in the presence of sub-lethal Hb. Previous research suggested that Hb can alter membrane potential [50], however the data was collected at a significantly higher concentration of 0.1 mM Hb, equivalent to 200 μM Hb dimer – a concentration that induced significant loss of ATP in our cultures. We intended to study the effects of one week exposure to sub-lethal Hb, hence we applied HL at 10 μM which did not result in an ATP deficit in our system and measured intrinsic membrane properties. We found no change to the resting membrane potential, input resistance or ability and frequency of firing action potentials when HL was applied for up to one week, with or without Hp. This data suggests that cultured neurons can maintain their membrane properties in the presence of 10 μM of free Hb. Maintaining a resting membrane potential around -70 mV in neurons is an active process, as ATP is required to power the sodium potassium exchange pump and preserve the correct electrochemical gradients across the cell membrane [70]. Input resistance is affected by a number of factors including the presence of K+ leak channels, open synaptic receptor channels and the surface area of cell membrane [71,72], and affects the excitability of the neuron [73,74]. Correct membrane potential and input resistance will affect the level of stimuli required to elicit an action potential, and are necessary to keep neurons at an optimal excitability within the neuronal network. These measures, in addition to rheobase and action potential number, were unchanged by 10 μM HL. We can conclude from this data that basal membrane properties and neuron excitability are normal in the presence of 10 μM HL, which correlates with a preservation of normal ATP levels under these conditions.

We also studied neurotransmission in neuron cultures, especially since prior evidence showed biochemical changes in synaptic composition [21,22,75] and cognitive impairment after SAH [25,26,76]. We focused on AMPA glutamate receptors, as much of the fast excitatory neurotransmission in the hippocampus is mediated by these tetrameric ionotropic receptors. Rapid depolarisation through AMPARs is typically required to enable action potential firing, following a summated postsynaptic response. We found a reduction in the amplitude of mEPSCs, evoked EPSPs and EPSCs after HL incubation, which provides strong evidence for a reduction in the number of AMPA receptors at the postsynaptic site of recorded neurons. Using Western blotting we also observed a reduction in GluA1 subunit expression in cell cultures, similar to previous research [75]. GluA1 is present in the majority of AMPA receptors in the hippocampus [77–79] and as such, a reduction in expression is likely to at least partially explain the reduction in amplitude of excitatory currents. This finding is similar to that previously seen in a mouse model of early Alzheimer’s disease [80], indicating shared features of AMPAR downregulation in neurodegeneration.

Previous research suggests that Hb can also potentiate the excitotoxic effects of other molecules, such as in the presence of excess neurotransmitters [81]. Glutamate-mediated neurotoxicity has been implicated in ischaemic and haemorrhagic stroke [53,82,83] and lesions in SAH follow the functional neuroanatomy rather than vascular architecture [83]. Iron levels after haemorrhage may play a role in this process by altering glutamate uptake [84]. Excess glutamate causing excitotoxicity can drive AMPAR internalisation and degradation leading to synaptic depression [70] in a homeostatic downscaling mechanism to preserve neuronal excitability [85].

Other contributing factors include alterations to receptor trafficking, which may link to the degree of neurite beading observed in the presence of free Hb. Degradation of microtubules within neurites will disrupt transport pathways within the cell, disturbing intracellular trafficking [86]. This could impair the trafficking of newly synthesised AMPA receptors to synaptic sites, which may have significant implications for synaptic plasticity. Hippocampal LTP requires the insertion of AMPA receptors into the synaptic membrane [87–89], and disruptions to trafficking via microtubules in addition to GluA1 protein downregulation will result in a lower availability of AMPA receptors available to enable long term potentiation mechanisms. These mechanisms may explain the loss of hippocampal LTP in a rat model of SAH [25], and since LTP is proposed to underpin many learning and memory processes [90], functional recovery after SAH is also likely impaired by changes to AMPAR availability.

The reduction in AMPAR-mediated currents observed in this study occurred at sublethal concentrations of free Hb which did not impair ATP or cause overt cell death, indicating that neurons in brain regions exposed to low levels of Hb may be surviving but are functionally impaired. Surviving cells exposed to Hb concentrations greater than 10 μM are likely to have even greater impairments in AMPA receptor-driven excitation, as evidence suggests that a reduction in ATP, as seen in our assays at 20 μM HL and above, leads to imbalance of Ca^2+^, Na^+^, Ca^2+^ and Cl^-^ ions, inhibiting glutamate reuptake [82]. However, we have not measured currents at greater HL concentrations using electrophysiology in this study, as visually guided patch clamp would be subject to selection bias of surviving cells.

We did not find any changes to presynaptic measures, including mEPSC frequency, evoked EPSP failure rate or paired-pulse facilitation. We also measured GABAA mIPSCs and found no change to frequency or amplitude (data not shown) suggesting sublethal Hb selectively impairs the postsynaptic element of AMPAR-mediated excitatory neurotransmission.

Throughout our data on AMPA receptor-mediated transmission deficits, we have seen that partially scavenging free Hb is sufficient to prevent a significant deficit. Hp binding of free Hb sequesters it in large stable complexes, preventing release of haem and compartmentalising the pathological species [91,92]. In this study it is likely that although we have only sequestered one-third of free Hb, this is sufficient to reduce a 10 μM concentration of Hb dimers to levels below the threshold for excitotoxicity or alteration to synapses; this is promising for the therapeutic potential of Hp.

It is also possible that Hp in itself has beneficial effects beyond binding Hb – for example Hp is well known as an anti-inflammatory protein, expressed in the acute phase response [93,94]. Since Hb breakdown products can activate inflammatory pathways in SAH [10,95,96] leading to activation of microglia and their conversion to harmful secretory phenotypes [9,97], Hp may provide neuroprotection beyond scavenging Hb through its anti-inflammatory effects. On its own, Hp did not impair ATP, and in fact a small increase in ATP levels was observed in neuronal cultures when incubated at 30 μM but no change from vehicle conditions was found at higher concentrations up to 120 μM. Neurite morphology was not altered by Hp, nor were intrinsic membrane properties or AMPA receptor-mediated excitatory currents. At the highest Hp concentration used in our study, Hp levels in culture medium were significantly higher than plasma concentrations [22] or those that would be delivered intrathecally for treatment of SAH, which is reassuring.

In conclusion, we isolated Hb neurotoxicity in a neuronal culture model of SAH and observed a dose-dependent impairment of ATP levels and neurite integrity upon long-term exposure to Hb. Clinically relevant, sub-lethal concentrations of Hb caused downregulation of AMPA receptors at the synapse, which may be due to a combination of altered protein expression levels and trafficking disruption from microtubule disintegration. Partially scavenging of one-third of free Hb using Hp was sufficient to restore AMPA and ATP deficits at 10-20 μM of HL, and Hp itself showed no negative effects even at very high concentrations. This data helps explain neurological deficits after SAH, and supports the development of Hp as a therapeutic agent to reduce the impact of secondary brain injury after SAH.

## List of abbreviations

AMPA: *α*-amino-3-hydroxy-5-methyl-4-isoxazoleproprionic acid
ATP: adenosine triphosphate
CSF: cerebrospinal fluid
DIV: day *in vitro*
EPSC: excitatory postsynaptic current
EPSP: excitatory postsynaptic potential
Hb: haemoglobin
Hp: haptoglobin
mEPSC: miniature excitatory postsynaptic current
RBC: red blood cell
SAH: subarachnoid haemorrhage
SBI: secondary brain injury.

## Declarations

### Ethics Approval

Human blood was used with consent (National research Ethics approval 12/SC/0176). Animal work was conducted in accordance with the Animals (Scientific Procedures) Act 1986 as approved by the UK Home Office.

## Consent for publication

Not applicable.

## Availability of data and material

The datasets used and/or analysed during the current study available from the corresponding authors on reasonable request.

## Authors’ contributions

HW, KD, DB, IG, and MVC designed experiments. H.W. performed experiments HW, KD and MVC and analysed the data. Haptoglobin was obtained and supplied by PG and JM All authors approved the manuscript.

## Acknowledgements

The authors thank Dr Charlotte Stuart for supply of human RBCs, and to Dr Mark Willett and the Imaging and Microscopy Centre at University of Southampton for training and support in this research.

## References

1. Petridis AK, Kamp MA, Cornelius JF, Beez T, Beseoglu K, Turowski B, et al. Aneurysmal subarachnoid hemorrhage-diagnosis and treatment. Dtsch. Arztebl. Int. Deutscher Arzte-Verlag GmbH; 2017. p. 226–35.

2. Hackett ML, Anderson CS. Health outcomes 1 year after subarachnoid hemorrhage an international population-based study. Neurology. Wolters Kluwer Health, Inc. on behalf of the American Academy of Neurology; 2000;55:658–62.

3. English SW. Long⍰Term Outcome and Economic Burden of Aneurysmal Subarachnoid Hemorrhage: Are we Only Seeing the Tip of the Iceberg? [Internet]. Neurocrit. Care. Springer; 2020. p. 37–8.

4. Luoma A, Reddy U. Acute management of aneurysmal subarachnoid haemorrhage. Contin Educ Anaesth Crit Care Pain. Narnia; 2013;13:52–8.

5. Diringer MN. Management of aneurysmal subarachnoid hemorrhage. Crit Care Med. NIH Public Access; 2009;37:432–40.

6. Diringer MN, Bleck TP, Hemphill JC, Menon D, Shutter L, Vespa P, et al. Critical care management of patients following aneurysmal subarachnoid hemorrhage: recommendations from the Neurocritical Care Society’s Multidisciplinary Consensus Conference. Neurocrit Care. Neurocrit Care; 2011;15:211–40.

7. Van Gijn J, Rinkel GJE. Subarachnoid haemorrhage: Diagnosis, causes and management. Brain. Oxford University Press; 2001. p. 249–78.

8. Yeh LH, Alayash AI. Redox side reactions of haemoglobin and cell signalling mechanisms. J Intern Med. John Wiley & Sons, Ltd; 2003. p. 518–26.

9. Kwon MS, Woo SK, Kurland DB, Yoon SH, Palmer AF, Banerjee U, et al. Methemoglobin is an endogenous Toll-like receptor 4 ligand—relevance to subarachnoid hemorrhage. Int J Mol Sci. MDPI AG; 2015;16:5028–46.

10. Belcher JD, Chen C, Nguyen J, Milbauer L, Abdulla F, Alayash AI, et al. Heme triggers TLR4 signaling leading to endothelial cell activation and vaso-occlusion in murine sickle cell disease. Blood. American Society of Hematology; 2014;123:377–90.

11. Joerk A, Seidel RA, Walter SG, Wiegand A, Kahnes M, Klopfleisch M, et al. Impact of heme and heme degradation products on vascular diameter in mouse visual cortex. J Am Heart Assoc. John Wiley and Sons Inc.; 2014;3.

12. Hugelshofer M, Buzzi RM, Schaer CA, Richter H, Akeret K, Anagnostakou V, et al. Haptoglobin administration into the subarachnoid space prevents hemoglobin-induced cerebral vasospasm. J Clin Invest. American Society for Clinical Investigation; 2019;129:5219.

13. Akeret K, Buzzi RM, Schaer CA, Thomson BR, Vallelian F, Wang S, et al. Cerebrospinal fluid hemoglobin drives subarachnoid hemorrhage-related secondary brain injury. J Cereb Blood Flow Metab. J Cereb Blood Flow Metab; 2021;41:3000–15.

14. Regan RF, Panter SS. Neurotoxicity of hemoglobin in cortical cell culture. Neurosci Lett. 1993;153:219–22.

15. Jungner Å, Vallius Kvist S, Romantsik O, Bruschettini M, Ekström C, Bendix I, et al. White Matter Brain Development after Exposure to Circulating Cell-Free Hemoglobin and Hyperoxia in a Rat Pup Model. Res Artic Dev Neurosci. 2019;41:234–46.

16. Wang X, Mori T, Sumii T, Lo EH. Hemoglobin-induced cytotoxicity in rat cerebral cortical neurons: Caspase activation and oxidative stress. Stroke. Lippincott Williams & Wilkins; 2002;33:1882–8.

17. Garton TP, He Y, Garton HJL, Keep RF, Xi G, Strahle JM. Hemoglobin-induced neuronal degeneration in the hippocampus after neonatal intraventricular hemorrhage. Brain Res. Elsevier; 2016;1635:86–94.

18. Zille M, Karuppagounder SS, Chen Y, Gough PJ, Bertin J, Finger J, et al. Neuronal Death after Hemorrhagic Stroke in Vitro and in Vivo Shares Features of Ferroptosis and Necroptosis. Stroke. Lippincott Williams and Wilkins; 2017;48:1033–43.

19. Li Q, Han X, Lan X, Gao Y, Wan J, Durham F, et al. Inhibition of neuronal ferroptosis protects hemorrhagic brain. JCI Insight. American Society for Clinical Investigation; 2017;2.

20. Bai Q, Liu J, Wang G. Ferroptosis, a Regulated Neuronal Cell Death Type After Intracerebral Hemorrhage. Front Cell Neurosci. Frontiers Media S.A.; 2020;14:374.

21. Shen H, Chen Z, Wang Y, Gao A, Li H, Cui Y, et al. Role of Neurexin-1β and Neuroligin-1 in Cognitive Dysfunction after Subarachnoid Hemorrhage in Rats. Stroke. Lippincott Williams & Wilkins Hagerstown, MD; 2015;46:2607–15.

22. Garland P, Morton MJ, Haskins W, Zolnourian A, Durnford A, Gaastra B, et al. Haemoglobin causes neuronal damage in vivo which is preventable by haptoglobin. Brain Commun. Oxford Academic; 2020;2.

23. Sasaki T, Hoffmann U, Kobayashi M, Sheng H, Ennaceur A, Lombard FW, et al. Long-Term Cognitive Deficits After Subarachnoid Hemorrhage in Rats. Neurocrit Care. Springer US; 2016;25:293–305.

24. Bliss TVP, Collingridge GL. A synaptic model of memory: long-term potentiation in the hippocampus. Nature. 1993;361:31–9.

25. Tariq A, Ai J, Chen G, Sabri M, Jeon H, Shang X, et al. Loss of long-term potentiation in the hippocampus after experimental subarachnoid hemorrhage in rats. Neuroscience. Pergamon; 2010;165:418–26.

26. Al-Khindi T, Macdonald RL, Schweizer TA. Cognitive and functional outcome after aneurysmal subarachnoid hemorrhage. Stroke. Lippincott Williams & Wilkins; 2010;41.

27. Tidswell P, Dias PS, Sagar HJ, Mayes AR, Battersby RDE. Cognitive outcome after aneurysm rupture: Relationship to aneurysm site and perioperative complications. Neurology. Wolters Kluwer Health, Inc. on behalf of the American Academy of Neurology; 1995;45:876–82.

28. Patel BN, Dunn RJ, Jeong SY, Zhu Q, Julien JP, David S. Ceruloplasmin regulates iron levels in the CNS and prevents free radical injury. J Neurosci. 2002;22:6578–86.

29. Núñez MT, Urrutia P, Mena N, Aguirre P, Tapia V, Salazar J. Iron toxicity in neurodegeneration. BioMetals. 2012;25:761–76.

30. Penke L, Valdés Hernandéz MC, Maniega SM, Gow AJ, Murray C, Starr JM, et al. Brain iron deposits are associated with general cognitive ability and cognitive aging. Neurobiol Aging. Elsevier; 2012;33:510-517.e2.

31. Jellinger KA. The role of iron in neurodegeneration. Prospects for pharmacotherapy of Parkinson’s disease. Drugs and Aging. 1999. p. 115–40.

32. Thomsen JH, Etzerodt A, Svendsen P, Moestrup SK. The haptoglobin-CD163-heme oxygenase-1 pathway for hemoglobin scavenging. Oxid Med Cell Longev. Hindawi Limited; 2013;2013:523652.

33. Kazmi N, Koda Y, Ndiaye NC, Visvikis-Siest S, Morton MJ, Gaunt TR, et al. Genetic determinants of circulating haptoglobin concentration. Clin Chim Acta. Elsevier; 2019;494:138–42.

34. Galea J, Cruickshank G, Teeling JL, Boche D, Garland P, Perry VH, et al. The intrathecal CD163-haptoglobin-hemoglobin scavenging system in subarachnoid hemorrhage. J Neurochem. Wiley-Blackwell; 2012;121:785–92.

35. Boretti FS, Buehler PW, D’Agnillo F, Kluge K, Glaus T, Butt OI, et al. Sequestration of extracellular hemoglobin within a haptoglobin complex decreases its hypertensive and oxidative effects in dogs and guinea pigs. J Clin Invest. American Society for Clinical Investigation; 2009;119:2271–80.

36. Gando S, Tedo I. The effects of massive transfusion and haptoglobin therapy on hemolysis in trauma patients. Surg Today. 1994;24:785–90.

37. Schaer DJ, Buehler PW, Alayash AI, Belcher JD, Vercellotti GM. Hemolysis and free hemoglobin revisited: exploring hemoglobin and hemin scavengers as a novel class of therapeutic proteins. Blood. 2013;121:1276–84.

38. Siddiqui FM, Bekker S V., Qureshi AI. Neuroimaging of Hemorrhage and Vascular Defects. Neurotherapeutics. Springer; 2011;8:28.

39. Janick PA, Hackney DB, Grossman RI, Asakura T. MR imaging of various oxidation states of intracellular and extracellular hemoglobin. Am J Neuroradiol. American Society of Neuroradiology; 1991;12:891–7.

40. Zhou J, Guo P, Guo Z, Sun X, Chen Y, Feng H. Fluid metabolic pathways after subarachnoid hemorrhage [Internet]. J. Neurochem. John Wiley & Sons, Ltd; 2022. p. 13–33.

41. Galea I, Durnford A, Glazier J, Mitchell S, Kohli S, Foulkes L, et al. Iron Deposition in the Brain after Aneurysmal Subarachnoid Hemorrhage. Stroke. Stroke; 2022;53:1633–42.

42. Galea I. The blood-brain barrier in systemic infection and inflammation. Cell Mol Immunol. Cell Mol Immunol; 2021;18:2489–501.

43. Nadkarni NA, Maas MB, Batra A, Kim M, Manno EM, Sorond FA, et al. Elevated Cerebrospinal Fluid Protein Is Associated with Unfavorable Functional Outcome in Spontaneous Subarachnoid Hemorrhage. J Stroke Cerebrovasc Dis. 2020;29:104605.

44. Benesch RE, Benesch R, Yung S. Equations for the spectrophotometric analysis of hemoglobin mixtures. Anal Biochem. Academic Press; 1973;55:245–8.

45. Schindelin J, Arganda-Carreras I, Frise E, Kaynig V, Longair M, Pietzsch T, et al. Fiji: an open-source platform for biological-image analysis. Nat Methods 2012 97. Nature Publishing Group; 2012;9:676–82.

46. Gibb AJ, Edwards FA. Patch clamp recording from cells in sliced tissues. Microelectrode Tech Plymouth Work Handb. 1994;255–74.

47. Schilling T, Eder C. Microglial K+ channel expression in young adult and aged mice. Glia. John Wiley & Sons, Ltd; 2015;63:664–72.

48. Stewart RR. Membrane properties of microglial cells isolated from the leech central nervous system. Proc R Soc London Ser B Biol Sci. The Royal SocietyLondon; 1994;255:201– 8.

49. Winchester G, Liu S, Steele OG, Aziz W, Penn A. Eventer. Software for the detection of spontaneous synaptic events measured by electrophysiology or imaging. 2020;

50. Yip S, Ip JKH, Sastry BR. Electrophysiological actions of hemoglobin on rat hippocampal CA1 pyramidal neurons. Brain Res. Elsevier; 1996;713:134–42.

51. Budohoski KP, Guilfoyle M, Helmy A, Huuskonen T, Czosnyka M, Kirollos R, et al. The pathophysiology and treatment of delayed cerebral ischaemia following subarachnoid haemorrhage [Internet]. J. Neurol. Neurosurg. Psychiatry. BMJ Publishing Group Ltd; 2014. p. 1343–53.

52. Neifert SN, Chapman EK, Martini ML, Shuman WH, Schupper AJ, Oermann EK, et al. Aneurysmal Subarachnoid Hemorrhage: the Last Decade [Internet]. Transl. Stroke Res. Springer; 2021. p. 428–46.

53. Righy C, T. Bozza M, F. Oliveira M, A. Bozza F. Molecular, Cellular and Clinical Aspects of Intracerebral Hemorrhage: Are the Enemies Within? Curr Neuropharmacol. 2015;14:392–402.

54. Cahill WJ, Calvert JH, Zhang JH. Mechanisms of early brain injury after subarachnoid hemorrhage [Internet]. J. Cereb. Blood Flow Metab. Nature Publishing Group; 2006. p. 1341–53.

55. Hugelshofer M, Sikorski CM, Seule M, Deuel J, Muroi CI, Seboek M, et al. Cell-Free Oxyhemoglobin in Cerebrospinal Fluid After Aneurysmal Subarachnoid Hemorrhage: Biomarker and Potential Therapeutic Target. World Neurosurg. Elsevier Inc.; 2018;120:e660–6.

56. Hilgenberg LGW, Smith MA. Preparation of Dissociated Mouse Cortical Neuron Cultures. J Vis Exp. 2007;

57. Hui CW, Zhang Y, Herrup K. Non-Neuronal Cells Are Required to Mediate the Effects of Neuroinflammation: Results from a Neuron-Enriched Culture System. PLoS One. Public Library of Science; 2016;11.

58. Regan RF, Guo Y. Toxic effect of hemoglobin on spinal cord neurons in culture. J Neurotrauma. Mary Ann Liebert Inc.; 1998;15:645–53.

59. Garland P, Broom LJ, Quraishe S, Dalton PD, Skipp P, Newman TA, et al. Soluble axoplasm enriched from injured CNS axons reveals the early modulation of the actin cytoskeleton. PLoS One. PLoS One; 2012;7.

60. Petzold A, Keir G, Kay A, Kerr M, Thompson EJ. Axonal damage and outcome in subarachnoid haemorrhage. J Neurol Neurosurg Psychiatry. BMJ Publishing Group Ltd; 2006;77:753–9.

61. Nylén K, Csajbok LZ, Öst M, Rashid A, Karlsson JE, Blennow K, et al. CSF –Neurofilament correlates with outcome after aneurysmal subarachnoid hemorrhage. Neurosci Lett. Elsevier; 2006;404:132–6.

62. Garland P, Morton M, Zolnourian A, Durnford A, Gaastra B, Toombs J, et al. Neurofilament light predicts neurological outcome after subarachnoid haemorrhage. Brain. Brain; 2021;144:761–8.

63. Gaastra B, Glazier J, Bulters D, Galea I. Haptoglobin Genotype and Outcome after Subarachnoid Haemorrhage: New Insights from a Meta-Analysis. Oxid Med Cell Longev. Hindawi; 2017;2017:1–9.

64. Gaastra B, Ren D, Alexander S, Bennett ER, Bielawski DM, Blackburn SL, et al. Haptoglobin genotype and aneurysmal subarachnoid hemorrhage individual patient data analysis. Neurology. Lippincott Williams and Wilkins; 2019;92:E2150–64.

65. Sadrzadeh SMH, Bozorgmehr J. Haptoglobin phenotypes in health and disorders. Am. J. Clin. Pathol. 2004.

66. Kwon SK, Kim MW. Pseudo-Froin’s syndrome, xanthochromia with high protein level of cerebrospinal fluid. Korean J Anesthesiol. Korean Society of Anesthesiologists; 2014;67:S58.

67. McCudden CR, Brooks J, Figurado P, Bourque PR. Cerebrospinal Fluid Total Protein Reference Intervals Derived from 20 Years of Patient Data. Clin Chem. Clin Chem; 2017;63:1856–65.

68. Jeffcote T, Ho KM. Associations between cerebrospinal fluid protein concentrations, serum albumin concentrations and intracranial pressure in neurotrauma and intracranial haemorrhage. Anaesth Intensive Care. Australian Society of Anaesthetists; 2010;38:274–9.

69. Chen-Roetling J, Regan RF. Haptoglobin increases the vulnerability of CD163-expressing neurons to hemoglobin. J Neurochem. Wiley/Blackwell (10.1111); 2016;139:586–95.

70. Zhang D, Hou Q, Wang M, Lin A, Jarzylo L, Navis A, et al. Na,K-ATPase Activity Regulates AMPA Receptor Turnover through Proteasome-Mediated Proteolysis. J Neurosci. Society for Neuroscience; 2009;29:4498–511.

71. Li Y-X, Zhang Y, Lester HA, Schuman EM, Davidson N. Enhancement of Neurotransmitter Release Induced by Brain-Derived Neurotrophic Factor in Cultured Hippocampal Neurons. 1998;

72. Breakdown S, Schmidt-hieber C, Jonas P, Bischofberger J, Jones P, Bischofberger J. Enhanced synaptic plasticity in newly generated granule cells of the adult hippocampus. Nature. Nature Publishing Group; 2004;429:184–7.

73. Kaczmarek LK, Jennings KR, Strumwasser F, Nairn AC, Walter U, Wilson FD, et al. Microinjection of catalytic subunit of cyclic AMP-dependent protein kinase enhances calcium action potentials of bag cell neurons in cell culture. Proc Natl Acad Sci. Proceedings of the National Academy of Sciences; 1980;77:7487–91.

74. Sun Z, Williams DJ, Xu B, Gogos JA. Altered function and maturation of primary cortical neurons from a 22q11.2 deletion mouse model of schizophrenia. Transl Psychiatry. Nature Publishing Group; 2018;8:1–14.

75. Han SM, Wan H, Kudo G, Foltz WD, Vines DC, Green DE, et al. Molecular alterations in the hippocampus after experimental subarachnoid hemorrhage. J Cereb Blood Flow Metab. SAGE Publications; 2014;34:108–17.

76. Ellmore TM, Rohlffs F, Khursheed F. FMRI of working memory impairment after recovery from subarachnoid hemorrhage. Front Neurol. Frontiers Media SA; 2013;4:179.

77. Schwenk J, Baehrens D, Haupt A, Bildl W, Boudkkazi S, Roeper J, et al. Regional Diversity and Developmental Dynamics of the AMPA-Receptor Proteome in the Mammalian Brain. Neuron. Cell Press; 2014;84:41–54.

78. Lu W, Shi Y, Jackson AC, Bjorgan K, During MJ, Sprengel R, et al. Subunit Composition of Synaptic AMPA Receptors Revealed by a Single-Cell Genetic Approach. Neuron. Elsevier Inc.; 2009;62:254–68.

79. Wenthold RJ, Petralia RS, Blahos J, Niedzielski AS, Blahos J II, Niedzielski AS. Evidence for multiple AMPA receptor complexes in hippocampal CA1/CA2 neurons. J Neurosci. Society for Neuroscience; 1996;16:1982–9.

80. Chang EH, Savage MJ, Flood DG, Thomas JM, Levy RB, Mahadomrongkul V, et al. AMPA receptor downscaling at the onset of Alzheimer’s disease pathology in double knockin mice. Proc Natl Acad Sci U S A. National Academy of Sciences; 2006;103:3410–5.

81. Regan RF, Scott Panter S. Hemoglobin potentiates excitotoxic injury in cortical cell culture. J Neurotrauma. Mary Ann Liebert Inc.; 1996;13:223–31.

82. Nagy Z, Nardai S. Cerebral ischemia/repefusion injury: From bench space to bedside. Brain Res. Bull. Elsevier; 2017. p. 30–7.

83. Schatlo B, Dreier JP, Gläsker S, Fathi AR, Moncrief T, Oldfield EH, et al. Report of selective cortical infarcts in the primate clot model of vasospasm after subarachnoid hemorrhage. Neurosurgery. 2010;67:721–8.

84. Yu J, Guo Y, Sun M, Li B, Zhang Y, Li C. Iron is a potential key mediator of glutamate excitotoxicity in spinal cord motor neurons. Brain Res. Elsevier; 2009;1257:102–7.

85. Hou Q, Gilbert J, Man HY. Homeostatic regulation of AMPA receptor trafficking and degradation by light-controlled single-synaptic activation. Neuron. Cell Press; 2011;72:806–18.

86. Sadleir KR, Kandalepas PC, Buggia-Prévot V, Nicholson DA, Thinakaran G, Vassar R. Presynaptic dystrophic neurites surrounding amyloid plaques are sites of microtubule disruption, BACE1 elevation, and increased Aβ generation in Alzheimer’s disease. Acta Neuropathol. Springer Verlag; 2016;132:235–56.

87. Kessels HW, Malinow R. Synaptic AMPA Receptor Plasticity and Behavior. Neuron. Cell Press; 2009. p. 340–50.

88. Benke T, Traynelis SF. AMPA-Type Glutamate Receptor Conductance Changes and Plasticity: Still a Lot of Noise. Neurochem Res. Springer New York LLC; 2019;44:539–48.

89. Gu J, Lee CW, Fan Y, Komlos D, Tang X, Sun C, et al. ADF/cofilin-mediated actin dynamics regulate AMPA receptor trafficking during synaptic plasticity. Nat Neurosci. Nature Publishing Group; 2010;13:1208–15.

90. Bliss TVP, Collingridge GL. A synaptic model of memory: Long-term potentiation in the hippocampus. Nature. 1993. p. 31–9.

91. Schaer CA, Jeger V, Gentinetta T, Spahn DR, Vallelian F, Rudiger A, et al. Haptoglobin treatment prevents cell-free hemoglobin exacerbated mortality in experimental rat sepsis. Intensive Care Med. Exp. SpringerOpen; 2021. p. 1–4.

92. Buehler PW, Humar R, Schaer DJ. Haptoglobin Therapeutics and Compartmentalization of Cell-Free Hemoglobin Toxicity. Trends Mol Med. Elsevier Current Trends; 2020;26:683– 97.

93. Moestrup S, Møller H. CD163: a regulated hemoglobin scavenger receptor with a role in the anti-inflammatory response. Ann Med. Taylor & Francis; 2004;36:347–54.

94. Landis RC, Philippidis P, Domin J, Boyle JJ, Haskard DO. Haptoglobin Genotype-Dependent Anti-Inflammatory Signaling in CD163(+) Macrophages. Int J Inflam. Hindawi; 2013;2013:980327.

95. Rosi MC, Luccarini I, Grossi C, Fiorentini A, Spillantini MG, Prisco A, et al. * Department of Preclinical and Clinical Pharmacology, University of Florence, Florence, Italy Department of Clinical Neurosciences, Brain Repair Centre, University of Cambridge, Cambridge, UK à Istituto di Genetica e Biofisica ‘“A. Buzzati Traverso”‘, CN. J Neurochem. 2010;112:1539–51.

96. Fassbender K, Hodapp B, Rossol S, Bertsch T, Schmeck J, Schütt S, et al. Inflammatory cytokines in subarachnoid haemorrhage: association with abnormal blood flow velocities in basal cerebral arteries. J Neurol Neurosurg Psychiatry. BMJ Publishing Group Ltd; 2001;70:534–7.

97. You W, Wang Z, Li H, Shen H, Xu X, Jia G, et al. Inhibition of mammalian target of rapamycin attenuates early brain injury through modulating microglial polarization after experimental subarachnoid hemorrhage in rats. J Neurol Sci. Elsevier; 2016;367:224–31.

